# Genome mosaicism in field strains of *Mycoplasma bovis* as footprints of *in-host* horizontal chromosomal transfer

**DOI:** 10.1101/2021.07.12.452010

**Authors:** Ana García-Galán, Eric Baranowski, Marie-Claude Hygonenq, Mathilda Walch, Guillaume Croville, Christine Citti, Christian De la Fe, Laurent-Xavier Nouvel

## Abstract

Horizontal gene transfer was long thought to be marginal in *Mollicutes*, but the capacity of some of these wall-less bacteria to exchange large chromosomal regions has been recently documented. Mycoplasma chromosomal transfer (MCT) is an unconventional mechanism that relies on the presence of a functional integrative conjugative element (ICE) in at least one partner and involves the horizontal acquisition of small and large chromosomal fragments from any part of the donor genome, which results in progenies composed of an infinitive variety of mosaic genomes. The present study focuses on *Mycoplasma bovis*, an important pathogen of cattle responsible for major economic losses worldwide. By combining phylogenetic tree reconstructions and detailed comparative genome analyses of 36 isolates collected in Spain (2016-2018) we confirmed the mosaic nature of 16 field isolates and mapped chromosomal transfers exchanged between their hypothetical ancestors. This study provides evidence that MCT can take place in the field, most likely during co-infections by multiple strains. Because mobile genetic elements (MGEs) are classical contributors of genome plasticity, the presence of phages, insertion sequences (ISs) and ICEs was also investigated. Data revealed that these elements are widespread within the *M. bovis* species and evidenced classical horizontal transfer of phages and ICEs in addition to MCT. These events contribute to wide-genome diversity and reorganization within this species and may have a tremendous impact on diagnostic and disease control.

**IMPORTANCE:** *Mycoplasma bovis* is a major pathogen of cattle with significant detrimental economic and animal welfare on cattle rearing worldwide. Understanding the evolution and the adaptative potential of pathogenic mycoplasma species in the natural host is essential to combating them. In this study, we documented the occurrence of mycoplasma chromosomal transfer, an atypical mechanism of horizontal gene transfer, in field isolates of *M. bovis* that provide new insights into the evolution of this pathogenic species in their natural host. Despite these events are expected to occur at low frequency, their impact is accountable for genome-wide variety and reorganization within *M. bovis* species, which may compromise both diagnostic and disease control.

## INTRODUCTION

Horizontal gene transfer (HGT) has deeply influenced our perception of bacterial evolution. By allowing the exchange of genetic material between organisms that are not in a parent-offspring relationship, HGT endows bacteria with the ability to eliminate deleterious mutations, respond to rapid environmental changes, and explore new ecological niches (1). Until recently, HGT was thought to be marginal in mycoplasmas (class *Mollicutes*), a large group of wall-less bacteria often portrayed as minimal cells because of their reduced genomes (ca. 0.5 to 2.0 Mb) and limited metabolic pathways (2). Indeed, genome erosion, —a degenerative process resulting from successive losses of genetic materials—, was long considered as the only force driving the evolution of *Mollicutes* from their Gram-positive ancestors (3). Recent evidence of massive HGT in *Mycoplasma* and other genera of the class *Mollicutes*, such as *Ureaplasma* or *Spiroplasma*, have provided a new frame to understand the evolution of these minimal bacteria (4–11). HGT in ruminant mycoplasmas was first documented by comparative genomic studies that have revealed the exchanges of large chromosomal regions between phylogenetically remote species known to share the same ecological niche (6). These data were further supported by the identification of integrative conjugative elements (ICE) in the genome of several ruminant mycoplasma species and the demonstration that these mobile genetic elements (MGE) were able to confer conjugative properties to their recipient strains (12, 13). Mycoplasma ICEs belong to a new family of transmissible elements that encode the machinery for their self-excision, transmission by conjugation, and random integration into the genome of the recipient cell, where they replicate as part of the host chromosome (11). Remarkably, ICEs were also found to play a key role in mycoplasma chromosomal transfer (MCT), an unconventional mechanism of HGT that involves the horizontal acquisition of small and large chromosomal fragments originating from any part of the donor genome. This distributive process results in replacing several regions of the recipient chromosome at homologous sites (14) and offers these minimal bacteria a remarkable adaptive potential by generating a complex progeny consisting of an almost infinitive variety of mosaic genomes (15, 16). A broad distribution of ICEs is observed among ruminant mycoplasmas, with two types circulating within phylogenetically distant species (17). The widespread occurrence of ICEs in ruminant mycoplasmas together with their capacity to promote conjugation which in turn allow MCT, raises questions regarding the potential emergence of mosaic genomes in the field.

The present study focuses on *Mycoplasma bovis*, a cattle pathogen that induces major economic impacts on the global livestock industry and causes mastitis, arthritis, pneumonia, keratoconjunctivitis, otitis media and genital disorders (18, 19). The spread of *M. bovis* infection among animals, herds, regions or countries is usually associated with animal movements and the introduction of asymptomatic carriers, which are occasionally shedding the pathogen in milk, colostrum, nasal or genital secretions (18–20). A recent molecular typing of *M. bovis* strains circulating in Spain revealed two subtypes (STs), both being found in beef and dairy cattle, regardless of regions and clinical signs (21). Remarkably, several animals were found to be concomitantly infected by the two *M. bovis* genotypes revealing a favourable epidemiological context for MCT in this pathogenic species. This particular context led us to investigate the occurrence of mosaic genomes among *M. bovis* isolates collected in Spain between 2016 and 2018 by whole-genome sequencing. Phylogenetic tree reconstructions based on multilocus sequence typing (MLST) and whole-genome sequence- single nucleotide polymorphisms (WGS-SNPs) revealed several discrepancies pointing toward mosaic genomes. Detailed comparative genomic analyses further confirmed the mosaic nature of these genomes and precisely mapped chromosomal fragments that had been exchanged between their hypothetical ancestors.

## RESULTS

### *M. bovis* genotypes circulating in Spain are worldwide distributed

A coarse-grained image of *M. bovis* genotypes circulating in Spain has been generated by single-locus typing using *polC* sequences (21, 22). Here, isolates belonging to *polC-*ST2 (n = 16) and *polC*-ST3 (n = 20) were further sequenced by Illumina technology, characterized by two MLST schemes (23, 24) and WGS-SNPs genotyping (Table 1). These data were combined with those available for MLST-2 in the PubMLST database (https://pubmlst.org/organisms/mycoplasma-bovis) to study the global population structure of *M. bovis*. The minimum spanning tree (MST) generated failed to reveal any correlation between a geographic origin and a particular ST (Fig. 1). Indeed, most lineages, with more than one isolate, included members from different countries or even different continents. Several notable exceptions included STs only detected in Canada (e.g. ST65), Israel (e.g. ST33), USA (e.g. ST15), China (e.g. ST90) and Japan (e.g. ST100). However, this apparent correlation is weak considering that those countries are the source of most isolates available in the database. In Spain, four of the seven STs identified, namely ST8, ST21, ST96 and ST122, have also been detected in other countries. These data suggest an efficient circulation of *M. bovis* between countries and even between continents.

**FIGURE 1.**
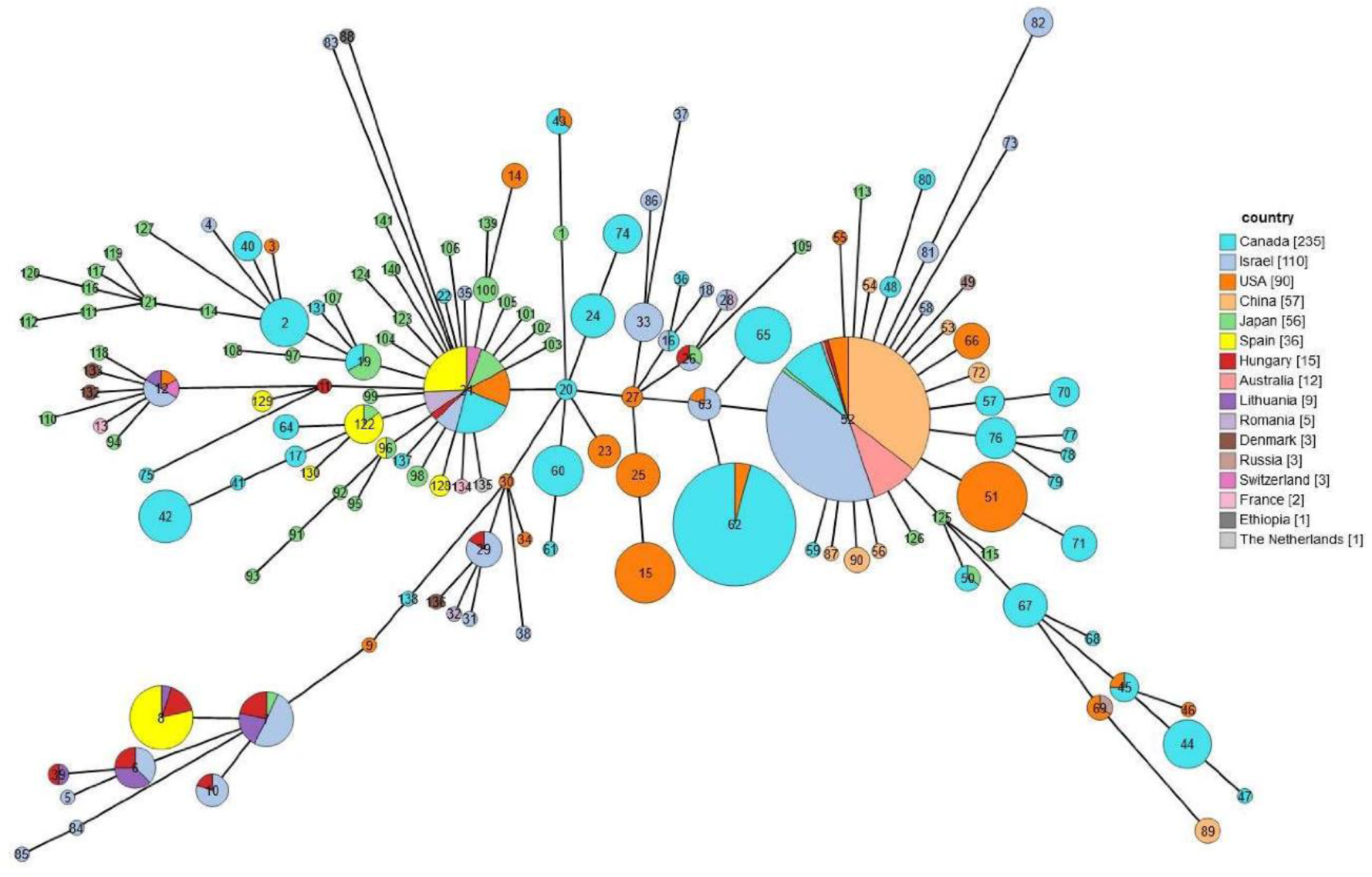
Global distribution of worldwide isolates using MLST-2 typing. MST representing the global distribution of the PubMLST STs (MLST-2) thus far defined for *M. bovis* until January 2020 and found among the isolates publically available in the database and with an assigned ST (https://pubmlst.org/organisms/mycoplasma-bovis). Each circle represents a unique ST indicated with a number. Circles or portions thereof, are sized in proportion to the number of isolates of each ST and are colored to indicate the country where they were found. In the legend, the number of isolates of each country is indicated in brackets. The 36 Spanish isolates are colored in yellow. The tree was created with GrapeTree (55).

**TABLE 1.**
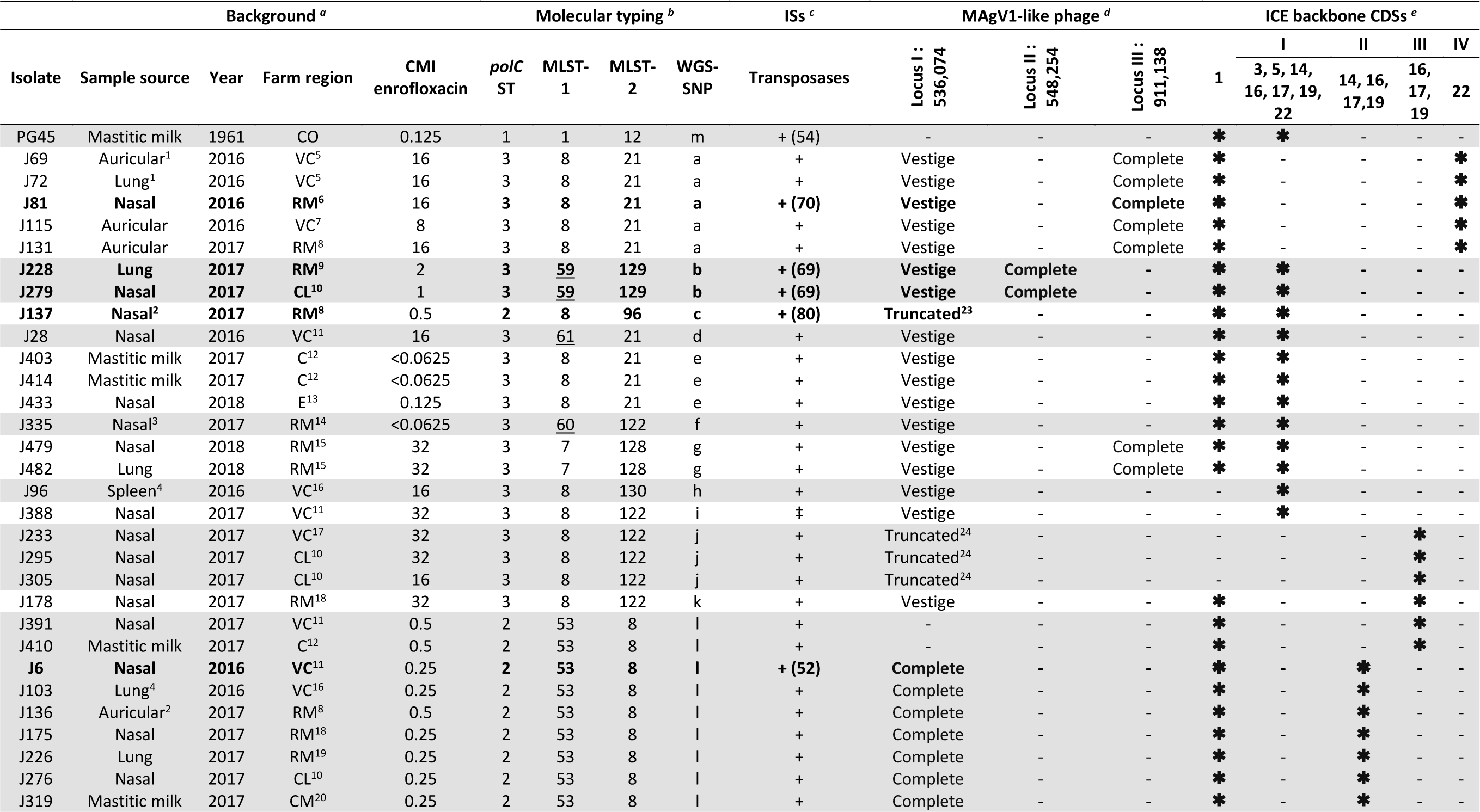
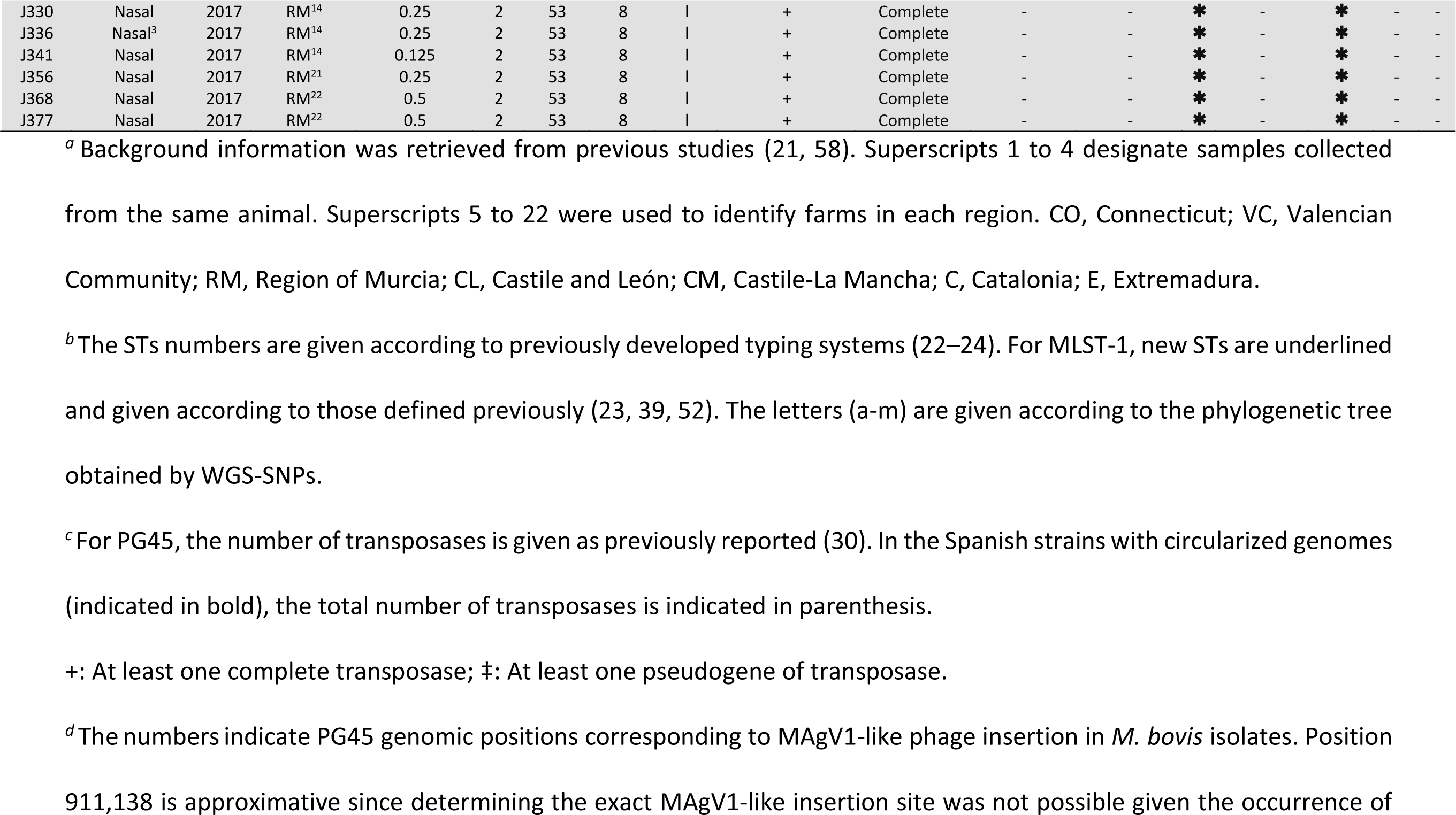
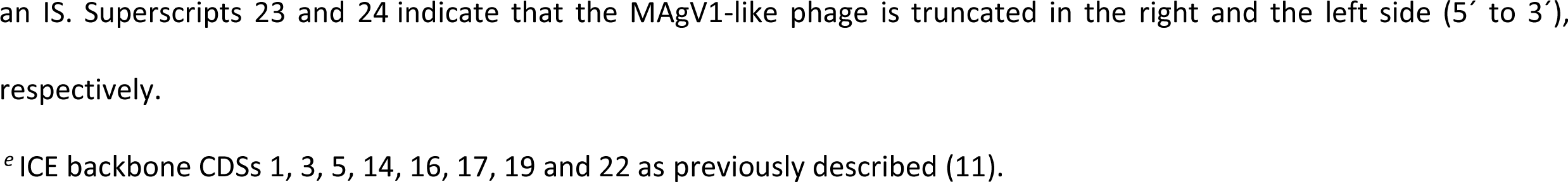
Epidemiological background, molecular characterization and mobilome composition of *M. bovis* isolates included in the study

### Identification of *M. bovis* isolates with a complex phylogenetic origin

Phylogenetic analyses based on selected loci versus whole genomic data revealed slightly different outcomes depending on the typing system (Table 1). MLST-1 identified seven STs, with the Spanish isolates clustering into three main branches (Table 1, Fig. S1). In contrast to most *polC-*ST2 isolates (15/16) that clustered into a single ST53, *polC*-ST3 isolates were found scattered into five STs and two main branches. One of these branches includes only two isolates (J228 and J279) belonging to ST59, which shares four alleles (*dnaA*, *recA*, *tufA* and *tkt*) with the reference strain PG45. Remarkably, one *polC-*ST2 isolate (J137) was classified into ST8, which mainly includes *polC*-ST3 isolates. Data generated with MLST-2 were consistent with MLST-1 (Table 1, Fig. 2), with only minor differences in the clustering of *polC-*ST3 isolates. These data confirm the proximity of J228 and J279 with PG45, which share four identical alleles (*dnaA*, *gltX*, *gyrB* and *tkt)*, as well as the clustering of J137 with *polC-* ST3 isolates. The phylogenetic analysis based on WGS-SNPs genotyping grouped Spanish isolates into two main branches: (i) branch A encompassing STs a to k, and (ii) branch B only composed of ST l (Table 1, Fig. 3). As expected, this system was highly discriminative, with up to 10 STs identified for *polC-*ST3 isolates in branch A, and two for *polC-*ST2 isolates, one of them being J137 (ST c) that clustered together with *polC-*ST3 isolates (Fig. 3). Finally, J228 and J279 were found to cluster in branch A, together with the other *polC*-ST3 isolates. No correlation was found between the occurrence of a particular ST and the sample source, year of isolation, or origin of the isolates in none of the three systems (Table 1). Overall, the three typing systems provide similar results showing two groups of isolates: one with a low diversity level (corresponding to *polC-*ST2 isolates) and one with higher heterogeneity (corresponding to *polC-*ST3 isolates). Remarkably, all molecular typing systems pointed towards three isolates, namely J137, J228 and J279, whose phylogenetic classification was inconsistent. Nanopore sequencing combined to Illumina permitted to obtain circularized genomes for these isolates (Fig. 4). Genome analyses further showed that J228 and J279 are nearly identical (Data set S1), with J279 having one additional CDS belonging to a family of lipoproteins, the DUF285 or PARCEL family, known to vary in gene number and expression within the species (25) when compared to J228. Below, we sometimes refer to these isolates as J228/279 keeping in mind this difference and that they were isolated from two distinct geographic areas in the same year.

**FIGURE 2.**
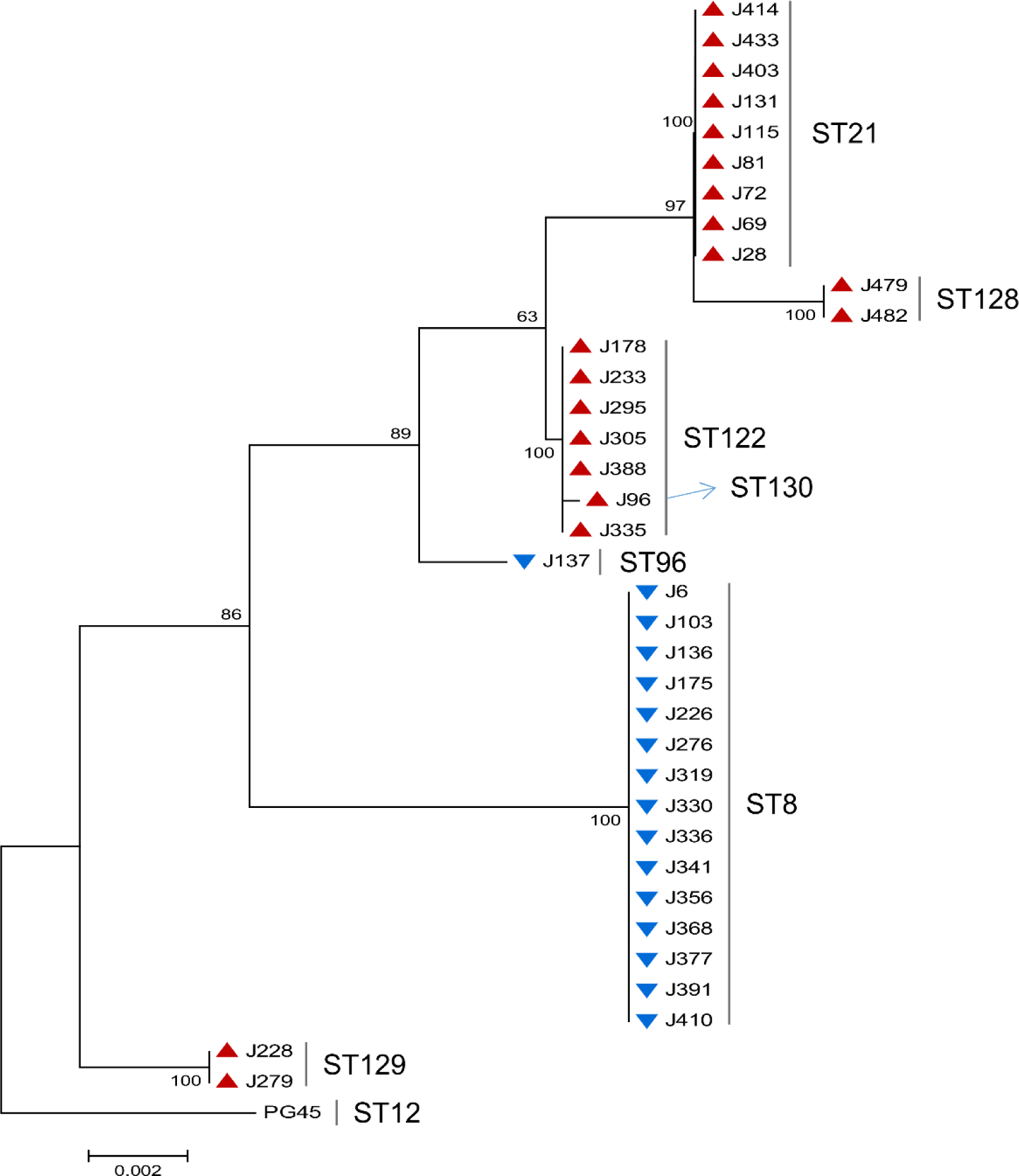
Molecular phylogenetic analysis based on MLST-2 concatenated sequences. Phylogenetic tree inferred from the concatenated partial sequences of the 7 genes included in the MLST-2 and corresponding to 36 *M. bovis* field isolates and the reference strain PG45 (CP002188.1). The red triangles refer to *polC*-ST3 isolates and the blue triangles to *polC*-ST2 isolates. The tree was constructed by using the Neighbor-Joining method, the Tamura-Nei genetic distance model, and 1000 bootstrap replicate analysis. Evolutionary analyses were conducted in MEGA X (51).

**FIGURE 3.**
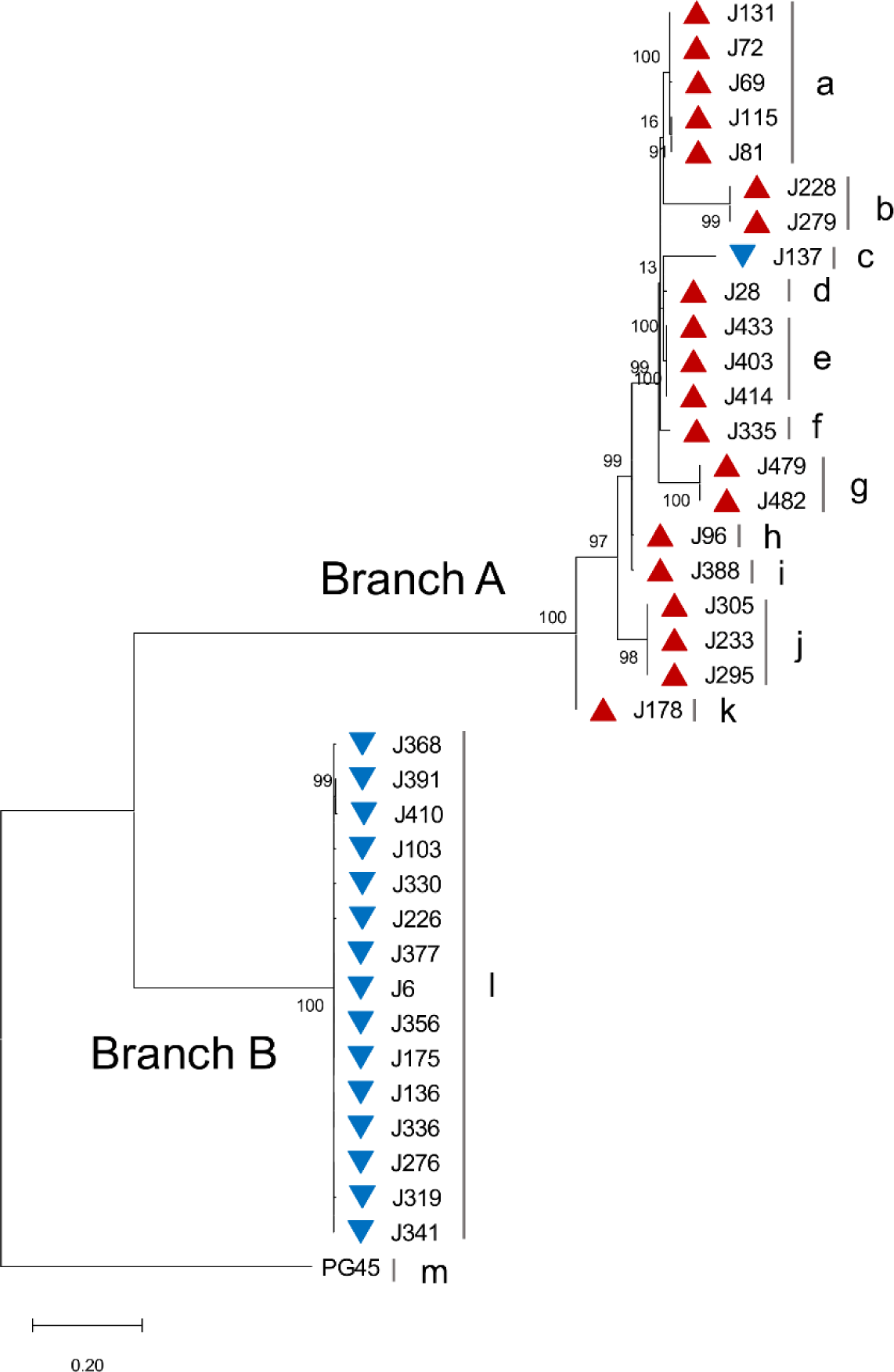
WGS-SNPs-based phylogenetic analysis of 36 *M. bovis* field isolates in comparison to the reference strain PG45. The red triangles refer to polC-ST3 isolates and the blue triangles to polC-ST2 isolates. The tree was constructed by using the Maximum Likelihood method, the Tamura-Nei genetic distance model, and 1000 bootstrap replicate analysis. Evolutionary analyses were conducted in MEGA X (51).

**FIGURE 4.**
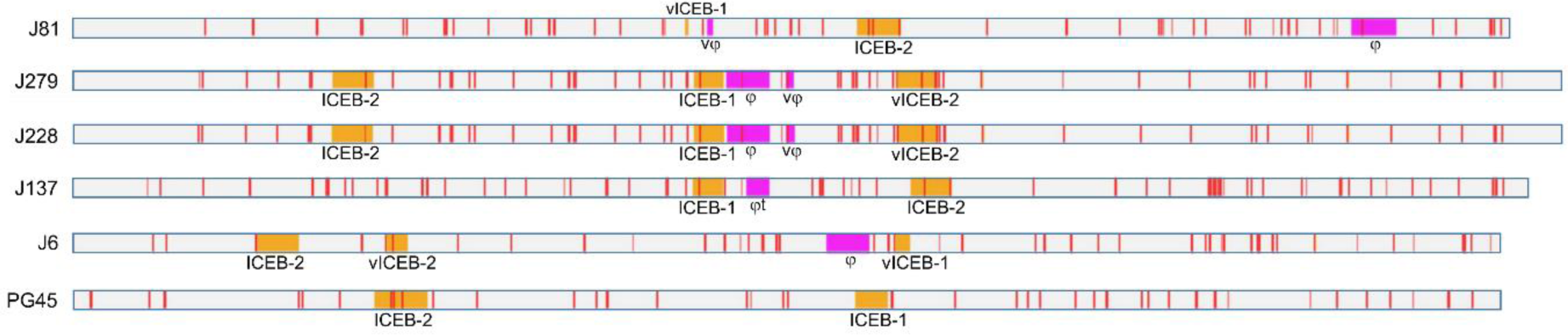
MGE genomic distribution among *M. bovis* genomes circularized in this study and reference strain PG45. DNAplotter linear representation of the genome circularized assemblies obtained with Oxford nanopore and Ilumina sequencing data (see Materials and Methods). ISs, MAgV1- like and ICEs are indicated in red, pink and orange respectively. ICE vestiges composed of less than two CDSs are not visualized but are described in Fig S3.

### MGEs are widespread among field strains

Because MGEs are known as the main contributors of genome plasticity, the presence of phages, ISs and ICEs was investigated in the set of 36 Spanish isolates. A MAgV1-like prophage, initially described in *M. agalactiae* (26) but absent from *M. bovis* type strain PG45, was recently identified in the French *M. bovis* isolate RM16 (27, 28). Homologous MAgV1-like sequences were found in all the Spanish strains except for two, J391 and J410, belonging to branch B (Table 1). More specifically, in branch B, prophage-positive strains all displayed a complete MAgV1-like version inserted at the same position (locus I, nucleotide 536,074 in PG45), while branch A strains only harbor vestigial or truncated prophage versions at this position (Table 1), with some (n = 9) displaying a complete prophage copy at a distinct locus (locus II, PG45 nucleotide 548,254; locus III, PG45 nucleotide 911,138). These findings, when superimposed to the phylogenetic history, suggest that branch A underwent two events: phage erosion leading to truncated or vestigial forms at locus 1 and independent episodes of phage attack at different loci (Table 1).

The overall genetic organization of the MAgV1-like prophage was compared in circularized genomes (J6, J81, J137, J228 and J279) using *M. bovis* RM16 as a reference (Fig. 5). The number of CDS, their orientation and size were highly similar to that found in RM16, the only differences being the occurrence of the IS element ISMbov4 in J81, J228 and J279. These strains also carried either prophage vestiges composed of a few CDSs (J81, J228 and J279) or truncated versions corresponding to about half of the prophage (J137) (Fig. 5).

**FIGURE 5.**
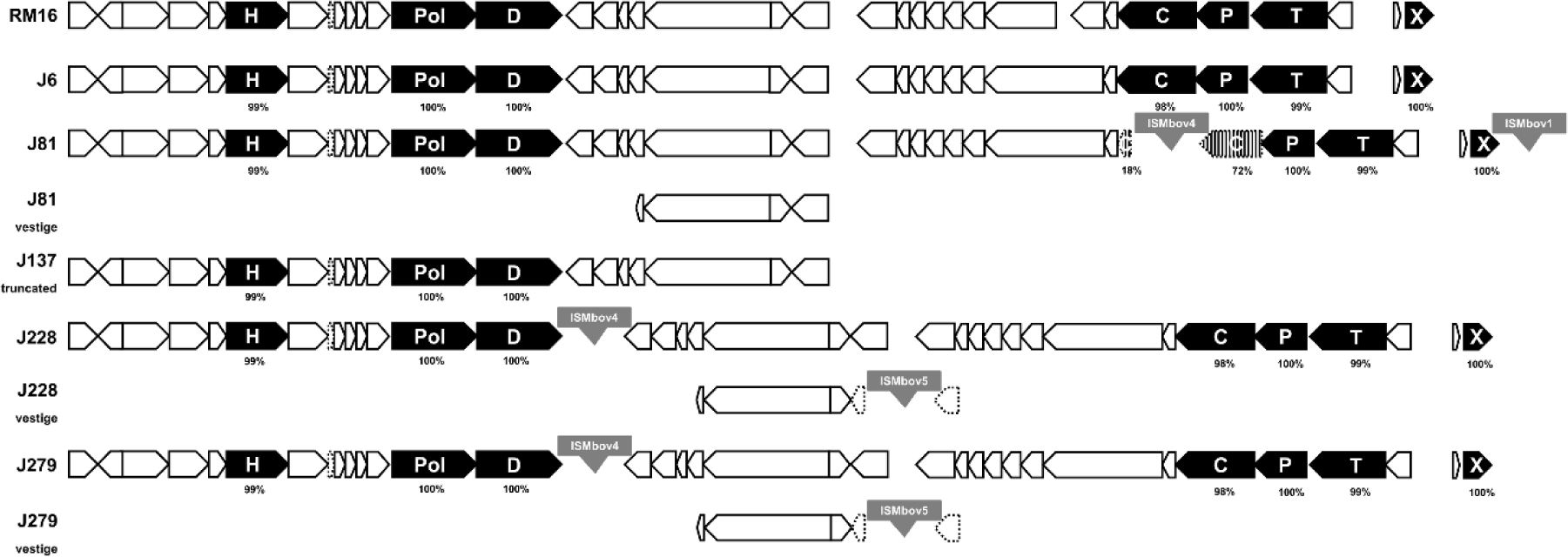
Comparison of complete, vestigial and truncated MagV1-like phages with that identified in *M. bovis* RM16 as reference (circularized in this study). The locations and orientations of the CDSs are indicated by arrows. CDSs highly conserved across MagV1-like phages identified across mycoplasma species are highlighted in black (26, 27). The letter code in black arrows refers to: H, helicase; Pol, DNA polymerase; D, DNA primase; C, prohead protein; P, portal; T, terminase; X, Xer. For each highlighted CDS, the percentage of global similarity with MagV1-like identified in RM16 was calculated by using the EMBOSS Needle alignment tool and it is indicated below (59). ISs are represented by grey boxes and are named according to the ISFinder database (29). CDSs interrupted by an IS are represented as pseudogenes with hatched colors and/or with dotted lines. MAgV1- like genomic positions and sizes are: RM16, 500818 to 532597 and 31.7 kb; J6, 548229 to 580006 and 31.7 kb; J81, complement (937518 to 970642) and 33.1 kb; J81vestige,complement (464793 to 469404) and 4.6 kb; J137_truncated_, complement (503457 to 521011) and 17.5 kb; J228, 500615 to 533739 and 33.1 kb; J228_vestige_, complement (546225 to 552494) and 6.2 kb; J279, 500599 to 533723 and 33.1 kb; J279_vestige_, complement (546209 to 552478) and 6.2 kb.

IS elements were identified in all Spanish isolates based on sequence similarities with transposases referenced in the ISFinder database (29) (Table 1). Between 52 to 80 transposases were identified in strains with circularized genomes (Table 1, Fig. 4). No correlation was found between the presence and/or the number of transposases and the epidemiological background or phylogenetic history of each strain.

Finally, particular attention was dedicated to ICEs, which are required for conjugation and MCT (11, 14). Their occurrence was determined by BLASTn using the two ICE sequences of the PG45 type strain, namely ICEB-1PG45 and ICEB-2PG45 (11, 30). Results showed that the Spanish isolates all possess at least one CDS of ICEB having a minimum coverage and nucleotide identity of 86% and 96%, respectively (Table 1). A minimal ICE backbone, composed of CDS1, CDS3, CDS5, CDS14, CDS16, CDS17, CDS19 and CDS22 as previously defined (11), was detected in 6 STs (b to g) that is 28 % (10/36) of the genomes. CDS1 was the most represented and was absent from only 5 isolates that clustered into three closely related STs, namely h, i, and j (Table 1). Overall, the isolates could be divided in 4 groups: one that perfectly matched ST a with isolates having only two CDSs, CDS1 and CDS22, and 3 others that displayed a combination of 3 (CDS16, 17 and 19), 4 (CDS14, 16, 17 and 19), or 7 CDS (CDS 3, 5, 14, 16, 17, 19 and 22), respectively, and with or without CDS1. Overall, the distribution of ICE CDSs was consistent with the phylogenic relationship of the strains and as expected there was no apparent relationship with their epidemiological background. Of the 5 isolates with a circularized genome, J81 (ST a), J228/279 (ST b), J137 (ST c) and J6 (ST l) (Fig. 4), only three, J137 and J228/279 had a complete ICEB-1 which organization was similar to that of ICEB-1PG45, except for an IS element interrupting CDS23 in both (Fig. S2). ICEB-2 also occurred in the circularized isolates (Fig. 4) but none that perfectly matched the reference ICEB-2PG45 (Fig S3) and both ICEB-1 and ICEB-2 were found in all isolates (Fig. 4). Of note, all ICEs forms detected in these isolates were associated with one or more ISs. Overall, MGEs, such as ICEs, MAgV1-like prophages and ISs, were widespread among *M. bovis* strains circulating in Spain (Table 1). There was no correlation between the distribution of these elements and the epidemiological background of the isolates. Data suggest that all these elements are active within the *M. bovis* species and thus constitute a source for genetic diversification.

### *M. bovis* isolates with a complex phylogenetic origin have a mosaic genome structure

The inconsistent phylogenetic allocation of J137 and J228/J279 suggested that these isolates may have a mosaic genome structure with some of their chromosomal regions originating from different parental isolates. To test this hypothesis, comparative genomic studies were performed with these 3 isolates and one representative genome of each main branch as potential parents. Isolates J81 and J6 were chosen as representative genomes of branches A and B respectively (Fig. 3) and were used to detect mosaic fragments in J137. Briefly, J137 reads were mapped onto the J81 genome, and those with mismatch were aligned to the J6 genome. Of these, reads perfectly matching the J6 genome were retained (Fig. 6, panel A). Data revealed that J137 displayed a J81 chromosomal backbone with 27 fragments (0.5-6.8 kb) identical to J6, which represent about 5.4 % of the genome (Table 2, Data set S2). One of these fragments included *polC,* explaining the inconsistencies observed with the phylogenetic trees based on this gene. As an internal control, J228 and J279 having similar genomic sequences (see above) but a slightly different sequence coverage (Table S1), were analyzed independently and shown to have a mosaic genome composed of a J81-backbone and 38 fragments identical to J6 encompassing about 7.1 % of each genome (Table 2). Using this strategy, genomes from branch A were further analyzed, and 13 additional mosaic genomes belonging to the phylogenetic STs d to k were identified. These genomes were characterized by a J81-backbone with 5 to 39 fragments identical to J6 (Table 2, Fig. 6 panel B). As expected, our approach did not detect any fragment potentially transferred in isolates of the phylogenetic ST a because of the parental reference J81 also belonging to this ST. Overall, up to 16 isolates of branch A were found to carry chromosomal fragments identical to J6 (branch B) and different from J81 (branch A). These mosaic genomes have acquired at least 5 to 39 fragments from a member or an ancestor of branch B which size ranged from 0.5 to 10.6 kb and represented about 0.9 to 8.5 % of the genomes (Table 2, Data set S2). As shown in Fig. 6 panel B, at least 8 different mosaic genomes were identified each corresponding to a different ST, except for d and e, and h and i that could not be distinguished. Close examination of transfered fragments showed that gene exchanges affected, for instance, several housekeeping genes like *gpsA*, *polC*, *gltX*, *pta-2*, *tufA* or *tkt* that were used for typing (22–24), as well as accessory putative lipoproteins and membrane proteins. Of note, fragments identified as being exchanged also included ICEs or part of ICEs. Whether these originated from chromosomal transfer or ICE self-transmission is not known as both events have been documented in previous studies using *M. agalactiae* (12, 14, 15).

**FIGURE 6.**
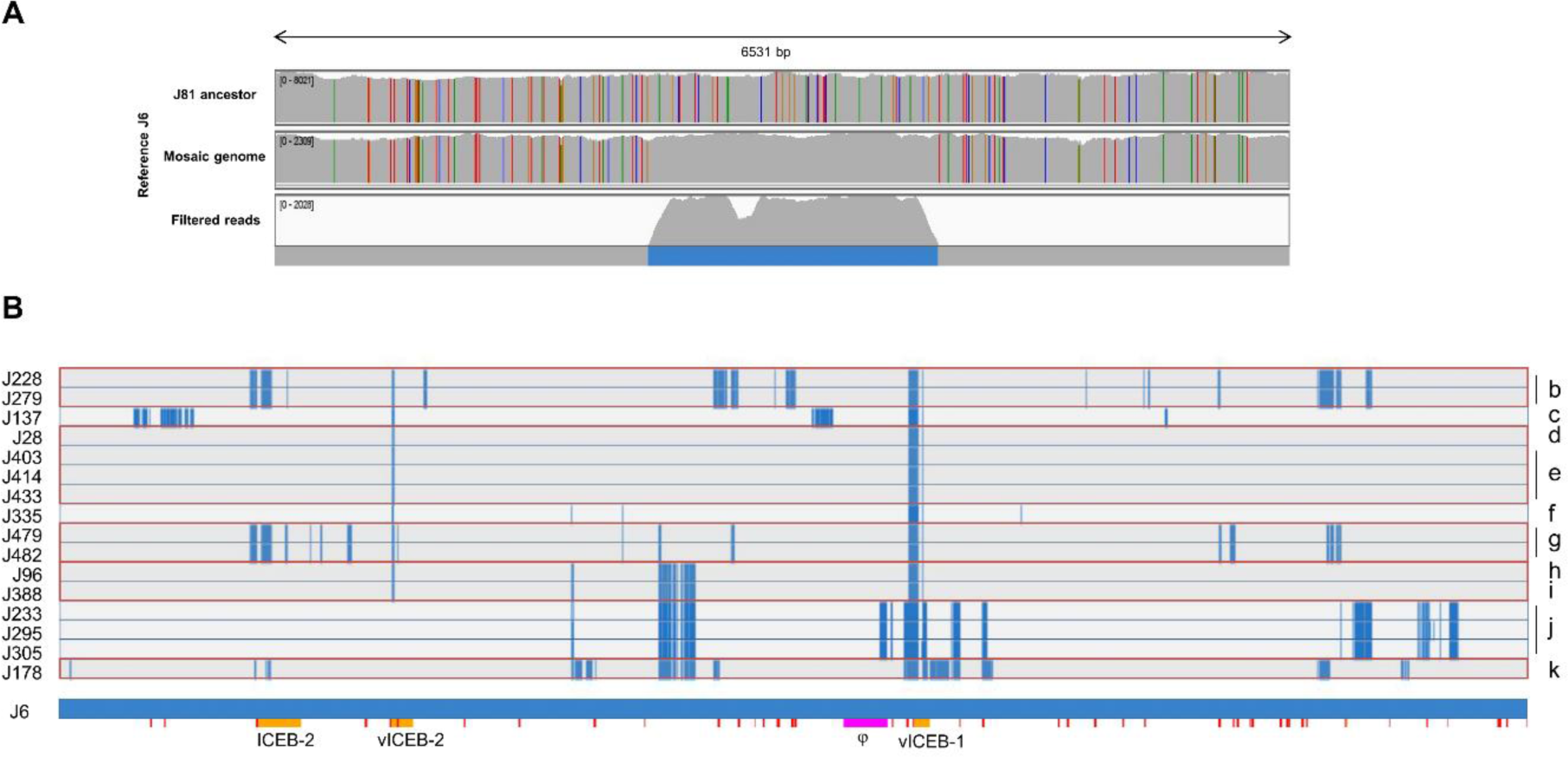
Mosaic genomes identified in *M. bovis* isolates of branch A. (A) Visualization of a transferred fragment in mycoplasma mosaic genomes using IGV 2.7.0 (57). Illumina reads were aligned to the J6 circularized genome (branch B), first line: branch A hypothetical ancestor (J81), second line: a mosaic genome (J137), third line: J137 reads filtered as different from J81 and identical to J6. Each color bar represents a SNP. The overall genotype of the mosaic genome is represented in the line below the alignments, with the genomic regions identical to J6 and J81 in blue and gray respectively. (B) Fragments of branch B origin identified in branch A isolates are represented: J6 (branch B) and J81 (branch A) specific sequences are indicated in blue and gray, respectively. ICEs and MAgV1-like relative positions in the J6 genome are indicated by orange and pink fragments respectively.

**TABLE 2.** Chromosomic fragments of branch A isolates potentially acquired from branch B

| Isolate | WGS-SNP<br>ST | Number of<br>fragments | % of genome<sup>a</sup> |
| --- | --- | --- | --- |
| J81 | a | - | - |
| J69 | a | 0 | 0 |
| J72 | a | 0 | 0 |
| J115 | a | 0 | 0 |
| J131 | a | 0 | 0 |
| J228 | b | 38 | 7.1 |
| J279 | b | 38 | 7.1 |
| J137 | c | 27 | 5.4 |
| J28 | d | 5 | 0.9 |
| J403 | e | 5 | 0.9 |
| J414 | e | 5 | 0.9 |
| J433 | e | 5 | 0.9 |
| J335 | f | 6 | 1.1 |
| J479 | g | 33 | 4.9 |
| J482 | g | 32 | 4.9 |
| J96 | h | 17 | 3.2 |
| J388 | i | 17 | 3.2 |
| J233 | j | 38 | 7.9 |
| J295 | j | 39 | 8.1 |
| J305 | j | 38 | 8 |
| J178 | k | 36 | 8.5 |

<sup>a</sup> the percentage was estimated by the total size of the fragments identical to J6 (branch B) and absent from J81 (branch A) divided by the size of the J6 genome (see Data set S2 for details of fragments positions for each isolate).

## DISCUSSION

By documenting the concomitant occurrence of two distinct mechanisms of HGT in field isolates of *M. bovis*, this study provides new insights into the evolution of pathogenic mycoplasma species in their natural host. Indeed, in addition to the classical horizontal transfer of MGEs, we provide evidence that MCT can take place in the field, most likely during co-infections by multiples strains. While these events are expected to occur at low frequency, their impact is responsible for genome-wide diversity and reorganization within the *M. bovis* species, which in turn may compromise both diagnostic and disease control, as further discussed.

MLST data indicated the circulation of multiple *M. bovis* STs in Spanish cattle from 2016 to 2018. Phylogenetic tree reconstruction based on WGS-SNPs genotyping further divided these isolates into two main branches, with branch A showing a greater level of intra-strain diversity than branch B. This was particularly true for MGEs, whose repertoire, distribution and location were more variable, as illustrated by MagV1-like prophage sequences. The MagV1 prophage was initially reported in the genome of atypical *M. agalactiae* strains associated with a mortality episode in Alpine ibexes in France (26). As evidenced in this study, this phage also circulates in *M. bovis* and the detection of a complete prophage sequence or vestigial forms at chromosomal position 536,074 in a majority of Spanish isolates (94 %) suggested an early viral attack, most likely in a common ancestor of the two branches. While a complete copy remained at this locus in most isolates of branch B, it was lost in two, collected the same year in distinct geographical areas. Whether these are originating from a single strain introduced in two distinct farms cannot be formally addressed, but their identical ICE pattern that differed from the rest of branch B argues in favor of this hypothesis. The situation was more complex in branch A where complete MagV1-like copies also occurred in addition to vestigial or truncated forms but at different chromosomal positions, either at nucleotide 548,254 or 911,138. Altogether, these data indicate that three independent genetic events have occurred after the splitting of the two branches. These are (i) the excision of the prophage resulting in virus-free isolates (branch B), (ii) a new viral attack that targeted two distinct loci (branch A), and (iii) the viral erosion or truncation of the copy vertically inherited (branch A). Remarkably, the MagV1-like prophage sequence appeared to be highly stable in *M. bovis*, with 99 % of nucleotide identity (over 97 % of the prophage sequence) in between the Spanish isolates and the Japanese strain KG4397 (AP019558). This suggests that the MagV1-like phage, which actively circulates in *M. bovis*, is under strong selection pressure. Whether this virus provides *M. bovis* with particular virulence or biological properties is unknown and many of its CDSs remain hypothetical, without any associated functions. Phages are not restricted to *M. bovis*, or its close relative *M. agalactiae*, but were detected in many other bovine mycoplasma species, such as *M. alkalescens, M. bovigenitalium, M. bovirhinis* and *M. arginini,* with the latter species being able to also infect a wide range of hosts in addition to ruminants (27).

Detailed genetic analyses also predicted the occurrence of single or multiple copies of entire and vestigial ICEs, with again more complex ICE patterns in branch A. Whether the entire ICEs described here are fully functional needs to be formally demonstrated but evidence for MCT supports this hypothesis. Indeed, mycoplasma ICEs are self-transmissible elements whose horizontal dissemination is not linked to MCT, yet MCT was shown to rely on the presence of functional ICE in at least one partner (11, 14). ISs were also highly widespread among contemporary Spanish isolates and contribute together with other MGEs, to genome plasticity by relocating themselves, disrupting genes or promoting genome re- arrangements (31). Hence, several ICEs and MAgV1-like CDSs were interrupted by ISs (Fig. 5, S2-S3) and major chromosome inversions were detected between strains J6 (branch B), J81 and J137 (branch A) that could be associated with MGE located nearby.

Discrepancies in phylogenetic tree reconstruction when using different typing methods raised the question of whether MCT could explain this result. MCT is an atypical, conjugative process of HGT documented for *M. agalactiae* in laboratory conditions. Because MCT involves any region of the genome that can be transferred alone or in combination, this phenomenon does not conform to canonical Hfr/*oriT* models of chromosomal transfer and results in progenies composed of a variety of mosaic genomes, each being unique. In-depth comparative genome analyses performed between a selection of Spanish *M. bovis* isolates of branch A, where a higher genetic diversity was observed, and one representative from each branch as potential parents revealed the circulation of several distinct mosaic genomes. More specifically, data showed the exchange of 5 to 39 fragments (0.9 to 8.5 % of the *M. bovis* genome) in between isolates of branches A and B. Our approach has several limits and these data might underestimate the total amount of DNA exchanges because it does not detect silent events resulting from the exchange of fragments with identical sequences. As well, the exact parents are not known and might even have not been collected in this study. Nevertheless, comparative genomic data were consistent with our MCT model (11) and thus strongly argue towards the occurrence of MCT events in *M. bovis* that have generated viable mosaic genomes currently circulating in the field.

Spanish isolates were associated with genomic STs already reported in various parts of the world suggesting a possible role for animal trading in the introduction of *M. bovis* strains in Spain, followed by an efficient transmission, circulation, and maintenance among and within the herds. Such a situation likely reflects animal movements between farms, a common practice in Spain (21). However, the genome mosaicism of several isolates also points towards MCT as an alternative cause for genetic diversity. When and where this phenomenon took place is unknown and, although this is probably not an isolated event, the frequency at which it occurs *in vivo* remains to be addressed. Interestingly, three animals turned out to be infected by two distinct isolates, one belonging to branch A, the other to branch B (Table 1). Since isolates from both branches were recovered from the same farms on several occasions, this raised the question of whether this observation is due to the independent introduction of two distinct isolates or whether it reflects the occurrence of MCT within the host, an exciting hypothesis that remains to be demonstrated. As illustrated here with MLST, MCT may have a tremendous impact on epidemiological studies. In mosaic genomes of branch A, the exchange of at least one housekeeping gene was detected including some used in *M. bovis-*specific MLST systems (22–24). These typing systems enable the characterization of *M. bovis* population structure and are used to trace the history and geographic route of dissemination of the pathogen. Hence, abrupt replacement of housekeeping genes by MCT could reduce the epidemiological value of MLST, as the variety of STs described in a given country, region or farm, may not only be the result of importing foreign strains but also the result of recombination between autochthonous strains. On the other hand, species-specific diagnosis may also be affected by MCT as many other mycoplasma species share the same ecological niche with *M. bovis* in cattle (14, 32, 33). Indeed, several studies pointed towards the ability of different species to engage in MCT. Indeed, MCT has been documented under laboratory conditions in between *M. bovis* and *M. agalactiae* (14), and *in silico* data showed that *M. agalactiae* has exchanged a large portion of its genome with phylogenetically remote ruminant mycoplasma species of the mycoides cluster (6). Thus MCT by altering the boundaries between species and strains may impact diagnostic in addition to epidemiological studies, a situation that may compromise the implementation of efficient control strategies. Putative membrane lipoproteins and membrane proteins were other transferred fragments. As mycoplasmas lack the cell wall, the membrane is in direct contact with the host and modification, gain or loss of this type of genes may drastically affect the host-pathogen interaction (34).

MCT may also compromise disease treatment. In *M. agalactiae,* MCT was shown to act as an accelerator of fluoroquinolone (FLQ) resistance dissemination *in vitro* under enrofloxacin selective pressure (16). Faucher *et al.* showed that MCT provided susceptible mycoplasma strains with the ability to rapidly acquire from pre-existing resistant populations, multiple chromosomal loci carrying antimicrobial mutations in *gyrA, parE* and *parC* genes. In Spain, circulating *M. bovis* isolates are resistant to macrolides, lincosamides and tetracyclines (21). Yet, while isolates in branch B were all susceptible to FLQ, most isolates found in branch A have acquired resistance-associated mutations. A single event of MCT, from FLQ-resistant to FLQ-susceptible isolates, followed by the expansion and transmission of the FLQ-resistant mosaic genome to other animals, farms or regions, may definitively compromise treatment of *M. bovis* infections with FLQ. Indirectly, animal movements between farms, a common practice in the Spanish beef cattle industry, would likely contribute to this situation by providing a means for mixing isolates with different antibiotic susceptibilities (21).

The high level of genetic diversity documented for branch A was consistent with previous observations showing a higher ability of *polC*-ST3 isolates to acquire mutations in the quinolone resistance-determining regions (QRDR) and develop resistance under selective pressure (35). The apparent selective advantage exhibited by branch A raises questions regarding the simultaneous circulation of another phylogenetic branch with limited genetic diversity. Other studies reported decreasing diversity and mono-clonal spread in several countries (22, 36–38). The existence of lineages with a high level of genetic diversity and others with low diversity has been described at worldwide scale (39, 40). This situation is not limited to *M. bovis* and has been observed in other mycoplasmas like in small ruminants with *M. agalactiae* (41, 42) where ovine isolates are described as more clonal whereas caprine are more variable (43). In *M. bovis* no correlation with host type or age was revealed. Our study highlights that the co-existence of lineages at the flock level and even at the animal level enables events of HGT like MCT. The occurrence, extension and evolution of this phenomenon have to be explored and integrated in control and diagnosis based on genomic typing.

## MATERIALS AND METHODS

### Mycoplasma sequences, genomes *de novo* assembly and annotation

Genome sequences of *M. bovis* reference strain PG45 (CP002188.1), RM16 and representative field isolates *polC-*ST2 (n = 16) and *polC*-ST3 (n = 20) collected between 2016 and 2018 in Spain were included in this study. The geographic and anatomic origins of the isolates at the time samples were collected are summarized in Table 1. These isolates were obtained in a previous study and sequenced using Illumina technology HiSeq (paired-end, 2x150 bp) (21). WGSs reads were retrieved from the European Nucleotide Archive database (ENA) (PRJEB38707). Bioinformatic analyses were conducted on the Galaxy Platform (Genotoul, Toulouse, France). The quality of the sequencing reads was analyzed with the FastQC tool (https://www.bioinformatics.babraham.ac.uk/projects/fastqc/). *De novo* assemblies were obtained with Unicycler (44). Five isolates sequenced by Illumina, identified as MCT potential parents or progenies, were also sequenced using the Oxford Nanopore Technology. RM16 strain (27, 28) was also re-sequenced with this technology. *M. bovis* genomic DNA was extracted using the phenol-chloroform method (45). Data were combined for genome assembly of these isolates to achieve chromosome circularization. For all genome assemblies obtained, contigs with less than 200 bp were excluded, validated with QUAST (46) and annotated with Prokka (47). Annotated files were visualized with Artemis 16.0.0 (48) and Artemis Comparison Tool (ACT 17.0.1) (49). Assembly metrics are available in Table S1. Average nucleotide identity (ANI) was calculated pairwise between assemblies of 36 Spanish isolates with FastANI-1.2 (50) (Data set S1).

### Phylogenetic isolates analysis

The phylogenetic relationships among the isolates were approached by tree reconstruction based on gene sequences of different MLST schemes and by WGS-SNP analyses.

Two MLST systems referred in the text to as MLST-1 (23) and MLST-2 (24) were used. Characterization with MLST-1 was based on the analysis of the partial sequence of *dnaA, metS, recA, tufA, atpA, rpoD, and tkt* genes (23). The amplicon sequences of these housekeeping genes were extracted from the annotated assemblies using Artemis 16.0.0 (48) and MLST primer sequences. They were trimmed to the analysis sequence size using MEGA X (51) and allelic profiles were assigned by blast comparison with the different allele sequences of each gene included in MLST-1 (23, 39, 52). New STs and allele sequences are provided in Tables S2 and S3. Typing with the MLST-2 was based on the analysis of the partial sequences of *dnaA*, *gltX*, *gpsA*, *gyrB*, *pta-2*, *tdk* and *tkt* (24). The STs were assigned with PubMLST database (automated sequence query for data derived from genome assemblies, https://pubmlst.org/bigsdb?db=pubmlst_mbovis_seqdef). New ST profiles were submitted to the database. Concatenated nucleotide sequences of each MLST system were used to reconstruct phylogenetic trees in MEGA X (51). The trees were constructed with the Neighbor-Joining method, the Tamura-Nei genetic distance model, and 1000 bootstrap replications.

The WGS-SNP-based analysis was performed by comparing the genome Illumina assemblies with PG45 genome (CP002188.1) as a reference on CSI phylogeny webserver (53) using parameters as previously described (54). Contigs of less than 1000 nucleotides were excluded. SNPs matrices generated by the server were used to create a phylogenetic tree using MEGA X (51). The tree was constructed with the Maximum Likelihood method, the Tamura-Nei genetic distance model, and 1000 bootstrap replications. A letter was manually assigned to each ST of isolates based on tree branches.

### Analysis of the global population structure of *M. bovis*

Minimum spanning tree (MST) that illustrates the STs distribution of isolates identified in this and other studies, was created with GrapeTree (55). Isolates registered with an assigned ST on PubMLST database (https://pubmlst.org/organisms/mycoplasma-bovis) (MLST-2) until January 2020 were included.

### Reconstruction of the mosaic genomes

The identification of strains with mosaic genomes and their hypothetical ancestors was based on the analysis of the phylogenetic trees. The short paired-end reads of possible isolates with mosaic genomes were mapped to the hypothetical ancestoŕs genomes by using BWA-MEM (56). The alignments were visualized with the Integrative Genome Viewer (IGV 2.7.0) (57). Reconstruction of the mosaic genomes was performed by “donor” specific reads detection as explained by Faucher et al (2019) (16) and Dordet-Frisoni et al (2019) (15): Reads of possible mosaic genomes were aligned on the “recipient” genome, reads with mismatch were recovered and then aligned to the “donor” genome. “Donor” specific reads were visualized in IGV 2.7.0 (57) and manually curated by using Artemis 16.0.0 and the Artemis BamView (48), and only reads with coverage higher than 50X and feature size higher than 500 bp were considered. False-positive fragments (also present in the negative control “recipient” ancestor), fragments present in the duplicated *rrn* operon and fragments having no SNPs based on Bam files were manually curated. Fragment plots were visualized with the Artemis 16.0.0 DNA plotter tool (48).

Additionally, the genome of three strains with mosaic genomes, namely J137, J228 and J279 was further circularized as described above. The effectiveness of the filtering process was then checked by mapping the paired-end reads of both hypothetical ancestors (“donor” and “recipient”) on the circularized genome of mosaic genomes and visualizing that, in the transferred regions, there were no SNPs in the alignment “donor”-mosaic genome but many SNPs in the alignment “recipient”-mosaic genome.

### Detection of MGE: ICEs, MAgV1-like phage and ISs

ICEs sequence detection was done by comparative analyses with the reference strain PG45 (CP002188.1), which carries two copies of ICE, ICEB-1 and ICEB-2 (30) . The ICE backbone CDSs sequences 3, 5, 14, 16, 17, 19, 22 of the ICEB-1 and the CDS1 sequence, which is only present in the ICEB-2, (GenBank locus-tags: MBOVPG45_0495 (CDS3), MBOVPG45_0494 (CDS5), MBOVPG45_0487 (CDS14), MBOVPG45_0484 (CDS16), MBOVPG45_0483 (CDS17), MBOVPG45_0481 (CDS19)

MBOVPG45_0479 (CDS22) and MBOVPG45_0213 (CDS1) were used as queries for BLASTn search against assemblies of the genomes studied.

Phage sequences were searched with MAgV1-like phage of *M. bovis* strain RM16 as reference, as it carries one copy of the phage (27, 28). Comparative analyses with PG45, which lacks the phage sequence, permitted to determine the phage insertion point. Finally, the presence of ISs was investigated by searches against the ISFinder database (29).

### Data availability

Nucleotide sequence assemblies were submitted to Genbank and are available under the following accession numbers: CP068734 (J6), JAERVI000000000 (J28), JAERVH000000000 (J69), JAERVG000000000 (J72), CP068733 (J81), JAERVF000000000 (J96), JAERVE000000000 (J103), JAERVD000000000 (J115), JAERVC000000000 (J131), JAERVB000000000 (J136), CP068732 (J137), JAERVA000000000 (J175), JAERUZ000000000 (J178), JAERUY000000000 (J226), CP068731 (J228), JAERUX000000000 (J233), JAERUW000000000 (J276), CP068730 (J279), JAERUV000000000 (J295), JAERUU000000000 (J305), JAERUT000000000 (J319), JAERUS000000000 (J330), JAERUR000000000 (J335), JAERUQ000000000 (J336), JAERUP000000000 (J341), JAERUO000000000 (J356), JAERUN000000000 (J368), JAERUM000000000 (J377), JAERUL000000000 (J388), JAERUK000000000 (J391), JAERUJ000000000 (J403), JAERUI000000000 (J410), JAERUH000000000 (J414), JAERUG000000000 (J433), JAERUF000000000 (J479), JAERUE000000000 (J482), CP077758 (RM16). Raw fastq reads used for mosaic genome reconstruction are available at the EMBL database, the European Nucleotide Archive (ENA) at http://www.ebi.ac.uk/ena under study accession number PRJEB38707.

## Supporting information

Pairwise average nucleotide identity (ANI) between assembly of 36 Spanish isolates

Detail of chromosomic fragments of branch A isolates potentially acquired from branch B

## ACKNOWLEDGMENTS

The authors are grateful to the Genotoul bioinformatics platform Toulouse Midi-Pyrenees for providing help and storage resources. This research has been supported by the Spanish Ministry of Economy and Competitiveness (Spanish Government) co-financed by FEDER funds (project AGL2016-76568-R), and financial supports from the INRAE and ENVT. Ana García-Galán Pérez is a beneficiary of a research fellowship (State Subprogram Training of the State Program for the Promotion of Talent and its Employability, BES-2017-080186). The funders had no role in study design, data collection and interpretation, or the decision to submit the work for publication.

