## Supplemental tables and figures for "Genome mosaicism in field strains of *Mycoplasma bovis* as footprints of *in-host* horizontal chromosomal transfer"

**SUPPLEMENTAL MATERIAL**

**Table S1.** Assembly metrics of the 36 *M. bovis* isolates included in this study

| Strain | Sequencing technology | Nº of contigs | Largest contig | Total length | Coverage | GC (%) | N50 |
|---|---|---|---|---|---|---|---|
| J6 | Illumina and Oxford Nanopore | 1 | 1038312 | 1038312 | 6679x | 29.34 | 1038312 |
| J28 | Illumina | 114 | 65669 | 941842 | 3221x | 29.32 | 19053 |
| J69 | Illumina | 106 | 70367 | 936676 | 8234x | 29.41 | 21684 |
| J72 | Illumina | 125 | 65791 | 949753 | 1755x | 29.42 | 21230 |
| J81 | Illumina and Oxford Nanopore | 1 | 1065959 | 1065959 | 7212x | 29.36 | 1065959 |
| J96 | Illumina | 104 | 64410 | 911129 | 3397x | 29.37 | 21681 |
| J103 | Illumina | 128 | 76097 | 935973 | 2738x | 29.37 | 21940 |
| J115 | Illumina | 121 | 65791 | 947790 | 2666x | 29.43 | 21387 |
| J131 | Illumina | 120 | 65791 | 948036 | 2158x | 29.41 | 21230 |
| J136 | Illumina | 109 | 59749 | 927408 | 1840x | 29.40 | 21940 |
| J137 | Illumina and Oxford Nanopore | 1 | 1088615 | 1088615 | 2216x | 29.20 | 1088615 |
| J175 | Illumina | 102 | 78827 | 924840 | 3137x | 29.40 | 22863 |
| J178 | Illumina | 105 | 49233 | 875841 | 3266x | 29.47 | 19180 |
| J226 | Illumina | 121 | 78828 | 937476 | 3274x | 29.39 | 22863 |
| J228 | Illumina and Oxford Nanopore | 1 | 1112468 | 1112468 | 2877x | 29.24 | 1112468 |
| J233 | Illumina | 112 | 65776 | 915159 | 1716x | 29.43 | 26400 |
| J276 | Illumina | 116 | 78827 | 924720 | 3066x | 29.39 | 22864 |
| J279 | Illumina and Oxford Nanopore | 1 | 1113982 | 1113982 | 2422x | 29.25 | 1113982 |
| J295 | Illumina | 113 | 65791 | 914730 | 3447x | 29.45 | 26836 |
| J305 | Illumina | 113 | 65777 | 913122 | 3988x | 29.44 | 26337 |
| J319 | Illumina | 114 | 78827 | 925487 | 5124x | 29.39 | 22863 |
| J330 | Illumina | 131 | 78822 | 936883 | 2807x | 29.39 | 23428 |
| J335 | Illumina | 109 | 65791 | 919598 | 4506x | 29.39 | 23165 |
| J336 | Illumina | 106 | 78827 | 925865 | 3858x | 29.40 | 23428 |
| J341 | Illumina | 116 | 78827 | 925162 | 4338x | 29.39 | 23428 |
| J356 | Illumina | 107 | 78827 | 924997 | 2078x | 29.39 | 23428 |
| J368 | Illumina | 115 | 78824 | 934837 | 2002x | 29.38 | 20736 |
| J377 | Illumina | 110 | 78827 | 924959 | 1978x | 29.40 | 23428 |
| J388 | Illumina | 107 | 65791 | 912866 | 4120x | 29.37 | 21230 |
| J391 | Illumina | 103 | 53371 | 893105 | 3849x | 29.36 | 21207 |
| J403 | Illumina | 129 | 43042 | 948447 | 3057x | 29.33 | 19182 |
| J410 | Illumina | 96 | 49485 | 892111 | 4397x | 29.35 | 21752 |
| J414 | Illumina | 132 | 43040 | 949474 | 2691x | 29.33 | 18099 |
| J433 | Illumina | 128 | 43040 | 948571 | 3057x | 29.33 | 19182 |
| J479 | Illumina | 137 | 70042 | 981092 | 2754x | 29.33 | 21366 |
| J482 | Illumina | 133 | 70042 | 979363 | 3333x | 29.34 | 21252 |
| RM16 | Illumina and Oxford Nanopore | 1 | 994478 | 994478 | 2000X | 29.50 | 994478 |

Contigs with size < 200 bp were excluded for the calculation of these values.

**Table S2.** New STs and allelic profiles identified in this study according to the MLST-1

| ST[a] | Allelic profile | | | | | | |
|---|---|---|---|---|---|---|---|
| | ***dnaA*** | ***metS*** | ***recA*** | ***tufA*** | ***atpA*** | ***rpoD*** | ***tkt*** |
| 59 | 1 | 5 | 1 | 1 | 2 | 5 | 1 |
| 60 | 1 | 5 | 1 | 2 | 2 | 13[b] | 4 |
| 61 | 1 | 5 | 1 | 13[b] | 2 | 5 | 4 |

[a] The ST number has been assigned considering the data previously published (1–3).

[b] New allele.

**Table S3.** New allele sequences of the *tufA* and *rpoD* genes according to the MLST-1

| > *tufA*_allele 13 |
|---|
| CGACCTTGTTGAAATGGAAGTTCGTGAATTACTTACAAAATATGGCTTTGATGGTGACAATACACCATTTGTTCGTGGTTCAGCTCTACAAGCTTTACAAA<br>GCAAACCAGAATACGAAGAAAACATTCTAGAATTAATGAATGCAGTTGACACATGAATCGAAACTCCTGTTAAAGACTTTGAAAAACCATTCTTAATGGC<br>TGTTGAAGACGTTTTCACAATTTCAGGTCGTGGTACAGTTGCTACAGGTCGTGTTGAACGTGGTAGATTAAGCTTAAACGAAGAAGTTGAAATTGTAGG<br>TCTTAAGCCAACAAAGAAAACAGTTGTTACAGGTATTGAAATGTTCAGAAAGAACTTAAAAGAAGCTCAAGCTGGTGACAACGCAGGTCTATTACTTCG<br>TGGTGTTGAAAGATCAGCTATTGAACGTGGACAAGTTCTTGCTAAACCAGGTTCTATTGTTCCTCATGCTGAATTCGAAGCAGCTATTTATGCATTAACA |
| > *rpoD*_allele 13 |
| TAGTGATAATTCTTATGATTCATACGAAGATGAATCTGAAGAGTTTATGGTTATGGATGATGATTACCATGACTCAAGCTATGACGAGGATGATGAAGA<br>CGACGAAATAGTCTATAAAACAGACGATAAAGACCAAGTCACTGCCAAAGCTAAAAGAGGAAGAAAACCTAAAACTAAACTTAATTTAGATACATTAGA<br>ATTAGAAGATTTCTCAGCTGAAGAAATTGATCTTAGTGCTAATTTATCAGTTAAAAAAGAAGATCTTAAAAACAAACTTACTGAAACAAATGATATTGTTA<br>AATGATACATGCGTTGAATTGGTAAGTATGGTAAATTATTGCAACCTGAAGAAGAATTTGAATTAGCTACTGAAATGGACAAAGGTGGTTTCAGAGGTA<br>AAAGAGCTAGAGATAAATTAATCAAAAGAAACTTGCGTTTGGTTATTAACAATGCTAAGAAGTACAAAAACAGGGGCTTAAGTTTTATTGACTTAATTTC<br>TGAAGGTAACTCAGGAATTATTAAGGCTGTGCAAAAGTACAATGTCAATAAAGGATTTAAATTTTCAACTTATGCAACTTGATGAATTCGTCAAGCAATT<br>AC |

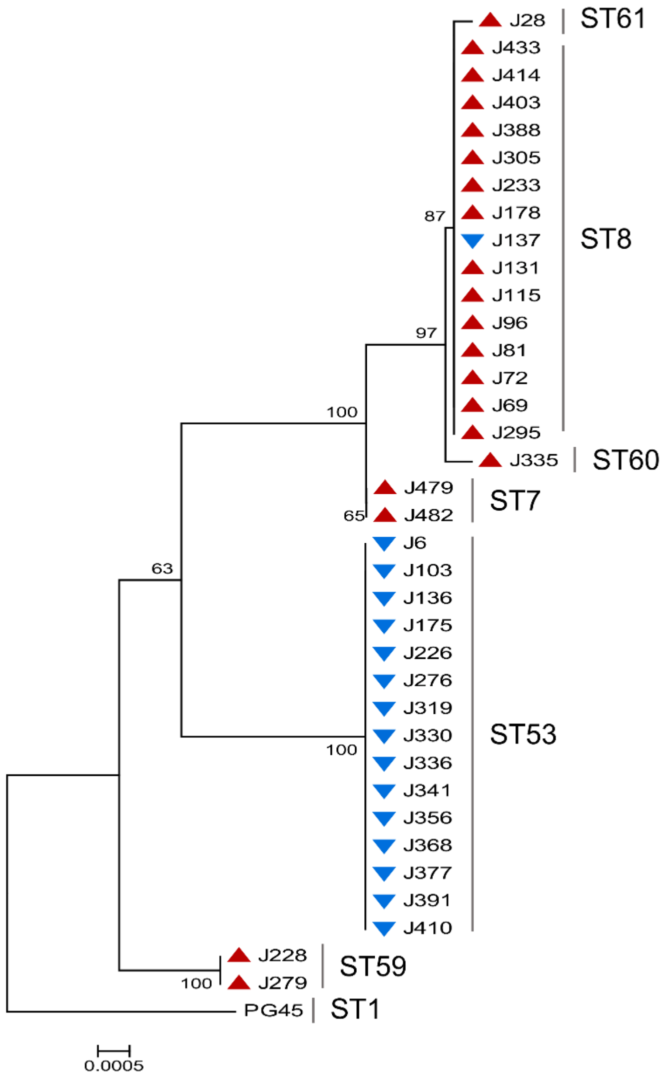


**FIGURE S1. Phylogenetic tree inferred from the concatenated partial sequences of the 7 genes included in the MLST-1 and corresponding to 36 *M. bovis* field isolates and the reference strain PG45 (CP002188.1).**

The red triangles refer to *polC*-ST3 isolates and the blue triangles to *polC*-ST2 isolates. The tree was constructed by using the Neighbor-Joining method, the Tamura-Nei genetic distance model, and 1000 bootstrap replicate analysis. Evolutionary analyses were conducted in MEGA X (4).

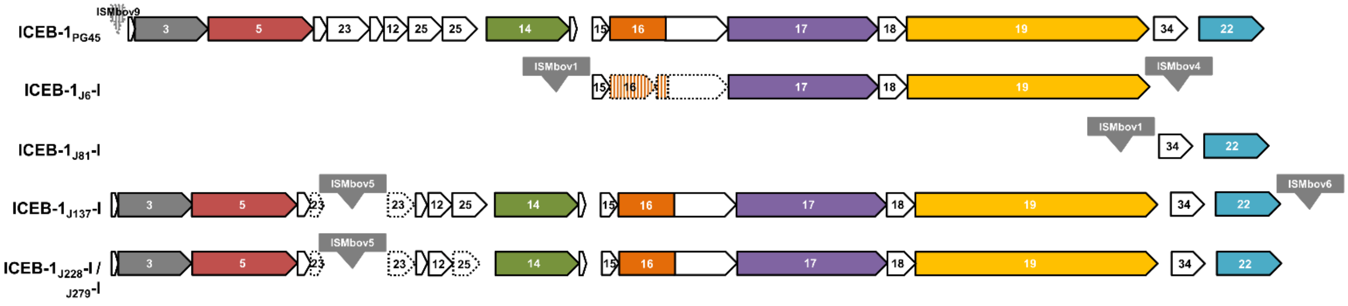


**FIGURE S2. Comparison of complete and vestigial ICEB-1 type ICEs with that identified in the PG45 reference strain (CP002188.1).**

The locations and orientations of the CDSs are indicated by arrows. The ICE backbone CDSs, which are those highly conserved across documented ICEs, are represented by colored filled arrows (5). ISs are represented by grey boxes and are named according to the ISFinder database (6). For each CDS, the percentage of global identity and similarity with its counterpart in PG45 was determined by using EMBOSS Needle alignment tool (7). CDSs with low conserved sequences (identity and similarity lower than 40 and 60%, respectively) were represented as pseudogenes by hatched colors and/or with dotted lines. Genes interrupted by an IS, or a premature stop codon truncating more than 16% of the product, were also represented as pseudogenes. ICEB-1 genomic positions and sizes are: ICEB-1$_{PG45}$, complement (550282 to 572181) and 21.9 kb; ICEB-1$_{J6}$-I, complement (598711 to 609336) and 10.6 kb; ICEB-1$_{J81}$-I, 448562 to 450757 and 2.1 kb; ICEB-1$_{J137}$-I, 463453 to 486470 and 23.0 kb; ICEB-1$_{J228}$-I, 475500 to 498519 and 23.0 kb; ICEB-1$_{J279}$-I, 475484 to 498503 and 23.0 Kb. For calculating the size of the ICE, the ISs positioned in the flanks were not considered.

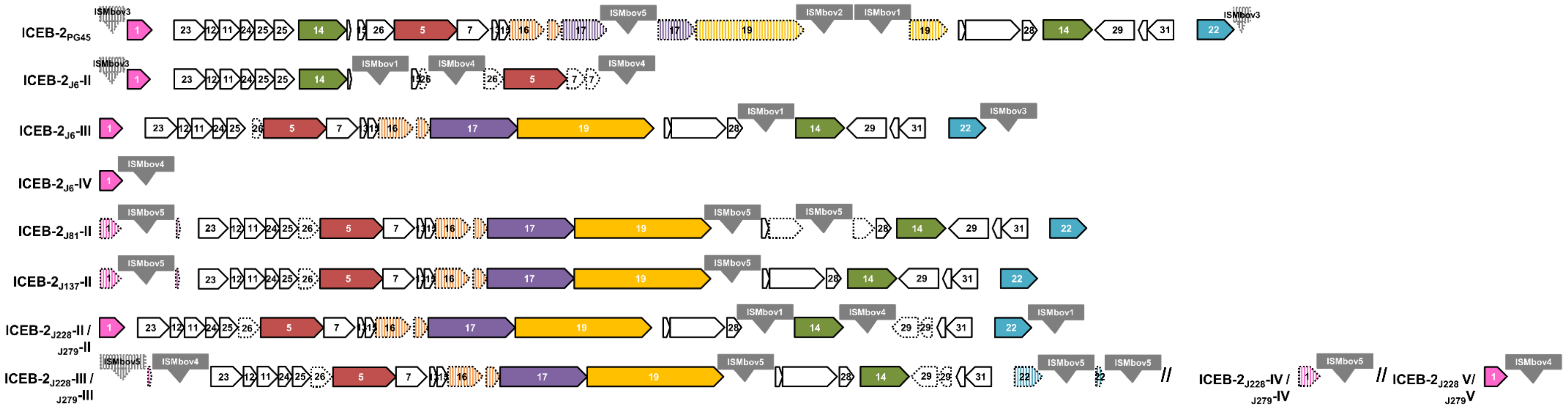


**FIGURE S3. Comparison of complete and vestigial ICEB-2 type ICEs with that identified in the PG45 reference strain (CP002188.1).**

The locations and orientations of the CDSs are indicated by arrows. ICE backbone CDSs, which are those highly conserved across documented ICEs, are represented by colored filled arrows (5). ISs are represented by grey boxes and are named according to the ISFinder database (6). For each CDS, the percentage of global identity and similarity with its counterpart in PG45 was determined by using EMBOSS Needle alignment tool (7). The CDSs 17 and 19 were aligned with their best blastp hit (BBH) with a conserved and characterized ICE CDSs, which was *M. agalactiae* 5362 strain (FP671138.1) in all the cases. CDSs with low conserved sequences (identity and similarity lower than 40 and 60%, respectively) were represented as pseudogenes by hatched colors and/or with dotted lines. Genes interrupted by an IS, or a premature stop codon truncating more than 16% of the product, were also represented as pseudogenes. MBOVPG45_0487, CDS14 carried by ICEB-1, is highly similar to MBOVPG45_0206, CDS14 carried by ICEB-2. ICEB-2 genomic positions and sizes are: ICEB-$2_{PG45}$, complement(210885..248018) and 37.1 kb; ICEB-$2_{J6}$-II, complement (228482 to 243652) and 15.1 kb; ICEB-$2_{J6}$-III, complement (134442 to 164288) and 29.8 kb; ICEB-$2_{J6}$-IV, complement (904169 to 904903) and 735 bp; ICEB-$2_{J81}$-II, complement (574673 to 607360) and 32.6 Kb; ICEB-$2_{J137}$-II, complement (626507 to 657536) and 31.0 kb; ICEB-$2_{J228}$-II, 198692 to 230111 and 31.4 kb; ICEB-$2_{J228}$-III, 630512 to 662240 and 31.7 kb; ICEB-$2_{J228}$-IV, complement (697116 to 697769) and 654 bp; ICEB-$2_{J228}$-V, complement (976069 to 976803) and 735 bp; ICEB-$2_{J279}$-II, 198678 to 230097 and 31.4 kb; ICEB-$2_{J279}$-III, 630495 to 662223 and 31.7 kb; ICEB-$2_{J279}$-IV, complement (697099 to 697752) and 654 bp; ICEB-$2_{J279}$-V, complement (977486 to 978220) and 735 bp. For calculating the size of the ICE, the ISs positioned in the flanks were not considered.
